# Using N2pc variability to probe functionality: Linear mixed modelling of trial EEG and behaviour

**DOI:** 10.1101/2024.05.31.596771

**Authors:** Clayton Hickey, Damiano Grignolio, Vinura Munasinghe, David Acunzo

**Author notes:** Correspondence to: Clayton Hickey, Centre for Human Brain Health University of Birmingham Edgbaston Birmingham, B60 2RU.

## Abstract

This paper has two concurrent goals. On one hand, we hope it will serve as a simple primer in the use of linear mixed modelling (LMM) for inferential statistical analysis of multimodal data. We describe how LMM can be easily adopted for the identification of trial-wise relationships between disparate measures and provide a brief cookbook for assessing the suitability of LMM in your analyses. On the other hand, this paper is an empirical report, probing how trial-wise variance in the N2pc, and specifically its sub-component the N_T_, can be predicted by manual reaction time (RT) and stimuli parameters. Extant work has identified a link between N2pc and RT that has been interpreted as evidence of a direct and causative relationship. However, results have left open the less-interesting possibility that the measures covary as a function of motivation or arousal. Using LMM, we demonstrate that the relationship only emerges when the N_T_ is elicited by targets, not distractors, suggesting a discrete and functional relationship. In other analyses, we find that the target-elicited N_T_ is sensitive to variance in distractor identity even when the distractor cannot itself elicit consistently lateralized brain activity. The N_T_ thus appears closely linked to attentional target processing, supporting the propagation of target-related information to response preparation and execution. At the same time, we find that this component is sensitive to distractor interference, which leaves open the possibility that N_T_ reflects brain activity responsible for the suppression of irrelevant distractor information.

## INTRODUCTION

Our visual environment contains enough information to overwhelm our cognitive capacity, and as a result we must prioritize information conveyed by some objects or locations at the expense of others. This is achieved in part through the deployment of spatial attention. When we attend to a lateral visual object, it has a neurophysiological correlate in the visual event-related-potential (ERP) that is known as the N2pc (Luck & Hillyard, 1994).

The N2pc emerges because of the lateralized organization of visual cortex. That is, objects or locations in the lateral periphery of the visual field are largely represented in the contralateral visual hemisphere. The deployment of attention to a lateral object causes an activity change in the neurons responsible for representation of this stimulus – thus in one visual cortex more than another – and this emerges in the ERP as a difference in voltage observed at electrodes located ipsilateral versus contralateral to the target. The component is evident in the contralateral-minus-ipsilateral difference wave at about 200 ms after stimulus onset and is named for its latency (in the range of bilateral N2) and *p*osterior *c*ontralateral scalp distribution.

Though evidence strongly indicates that the N2pc tracks attentional engagement, the precise mechanistic function of the underlying brain activity remains unclear. Seminal early work proposed that the component might reflect the suppression of distractors (eg. Luck & Hillyard, 1994b; Luck, Girelli, McDermott, & Ford, 1997), in line with contemporary results and theory from animal research (eg. Moran & Desimone, 1985; Chelazzi, Miller, Duncan, & Desimone, 1993). While this distractor suppression hypothesis remains viable, subsequent results suggest there are at least nuances to the account. That is, the N2pc appears under circumstances where it makes little sense to suppress distractors (Mazza, Turatto, & Caramazza, 2009). When displays contain only two visual objects in contralateral visual hemifields, it emerges contralateral to the target (Eimer, 1996), which is inconsistent with the common-sense notion that suppression of the distractor should have a neurophysiological correlate in cortex contralateral to the distractor itself. It can arguably be identified in response to single stimulus, presented in the absence of any distractors (Shedden & Nordgaard, 2001). And it appears to have distinct sub-components: the target negativity (Nt), which emerges contralateral to target stimuli and varies as a function of target features and location, and the distractor positivity (Pd), which emerges contralateral to distractors and appears to track their suppression (Hickey, Di Lollo, & McDonald, 2009; Weaver, van Zoest, & Hickey, 2017; Gaspelin et al., 2023).

In many ways, the identification of Pd has clarified the relationship between N2pc and direct suppression of distractor representation. But the functional role of N_T_ remains unclear. The fact that N_T_ is closely linked to target features and locations is in line with the broad proposal that N2pc reflects target processing (eg. Eimer & Kiss, 2008, 2010; Wyble et al., 2020). However, this idea does not have a clear definition in neurophysiological terms, at least to the level of specificity provided by the distractor suppression hypothesis. If the N_T_ reflects target processing, what exactly does this mean? One possibility deserving of consideration is that the N_T_ reflects neural activity involved in feature binding – more precisely, the reentrant boost of firing rates that is thought to underlie the establishment of cell assemblies to this end (Roelfsema, 2023; Di Lollo, 2012). But a valid alternative is that the N_T_ reflects the sheltering of target information through the inhibition of input from distractors into tissue responsible for target representation, and thus remains indirectly linked to distractor processing (Luck et al., 1997; Hickey, Di Lollo, & McDonald, 2009). This latter scenario would imply that attentional selection, as reflected in N2pc, depends on two distinct types of distractor suppression: one which acts directly on the cortical representation of distractors and is reflected in Pd, and one which acts to protect the target representation – perhaps through inhibitory laminar interactions (Priebe & Ferster, 2008) – and is reflected in N_T_.

This ambiguity in interpretation highlights the continuing need for investigation of the functional role of N2pc and its subcomponents. Here we pursue this via the identification of trial-wise covariance between N2pc and other trial-varying experimental parameters and measures, including reaction time (RT). The analysis of N2pc as a function of RT is not a new approach. In the past, N2pc results have been partitioned in this way to identify if attentional selection of highly salient distractors drives slowing in the behavioural response to targets (Hickey, van Zoest, & Theeuwes, 2010; McDonald, Green, Jannati, & Di Lollo, 2013). Building from this, Drisdelle, West, and Jolicoeur (2016) looked to the relationship between target-elicited N2pc and target RT in a visual search task. These authors conducted a median split analysis of within-participant results, generating separate ERPs for the set of trials associated with fast RT and the set of trials associated with slow RT. Results showed that the N2pc was larger in amplitude when responses were ultimately quicker.

Drisdelle, West, and Jolicoeur (2016) interpreted this as evidence of a direct relationship between the quality of attentional selection – represented in N2pc – and the ultimate speed of response. The logic was that when stimuli were better attended, the information they contained was quicker to reach response preparation and execution. This is compelling, but there is an alternative, namely that the relationship between N2pc and RT reflects the broad influence of motivation or arousal. According to this ‘third variable’ account, target-elicited N2pc and target RT are both independently sensitive to fluctuation in participant state, and it is this that creates covariance in the measures, rather than any more direct link.

A first goal for the current paper is therefore to clarify the relationship between N2pc and RT. Our experimental design (Figure 1) allowed us to isolate the N_T_ subcomponent elicited when attention was deployed to targets, but also to identify the N_T_ elicited by distractors when they were erroneously attended. We did this by presenting targets and distractors either at lateral locations – a circumstance where attentional selection would generate the N_T_ – or on the vertical meridian of the visual field – when selection will cause equal magnitude activity in both visual cortices and therefore no N_T_ (Woodman & Luck, 2003; Hickey, McDonald, & Theeuwes, 2006; Hickey et al., 2009). This allowed us to isolate the target-elicited N_T_ – evoked when the lateral target was attended – and the distractor-elicited N_T_ – evoked when the lateral distractor was attended. If N_T_ and RT broadly covary as a function of participant state, the relationship between these measures should emerge when target-elicited N_T_ is related to RT, but also when distractor-elicited N_T_ is related to RT. In contrast, if N_T_ has a relationship with RT through its impact on information processing and response preparation, the beneficial relationship between N_T_ and RT – wherein bigger N_T_ predicts faster RT – should only emerge in analysis of target-elicited N_T_.

**Figure 1.**
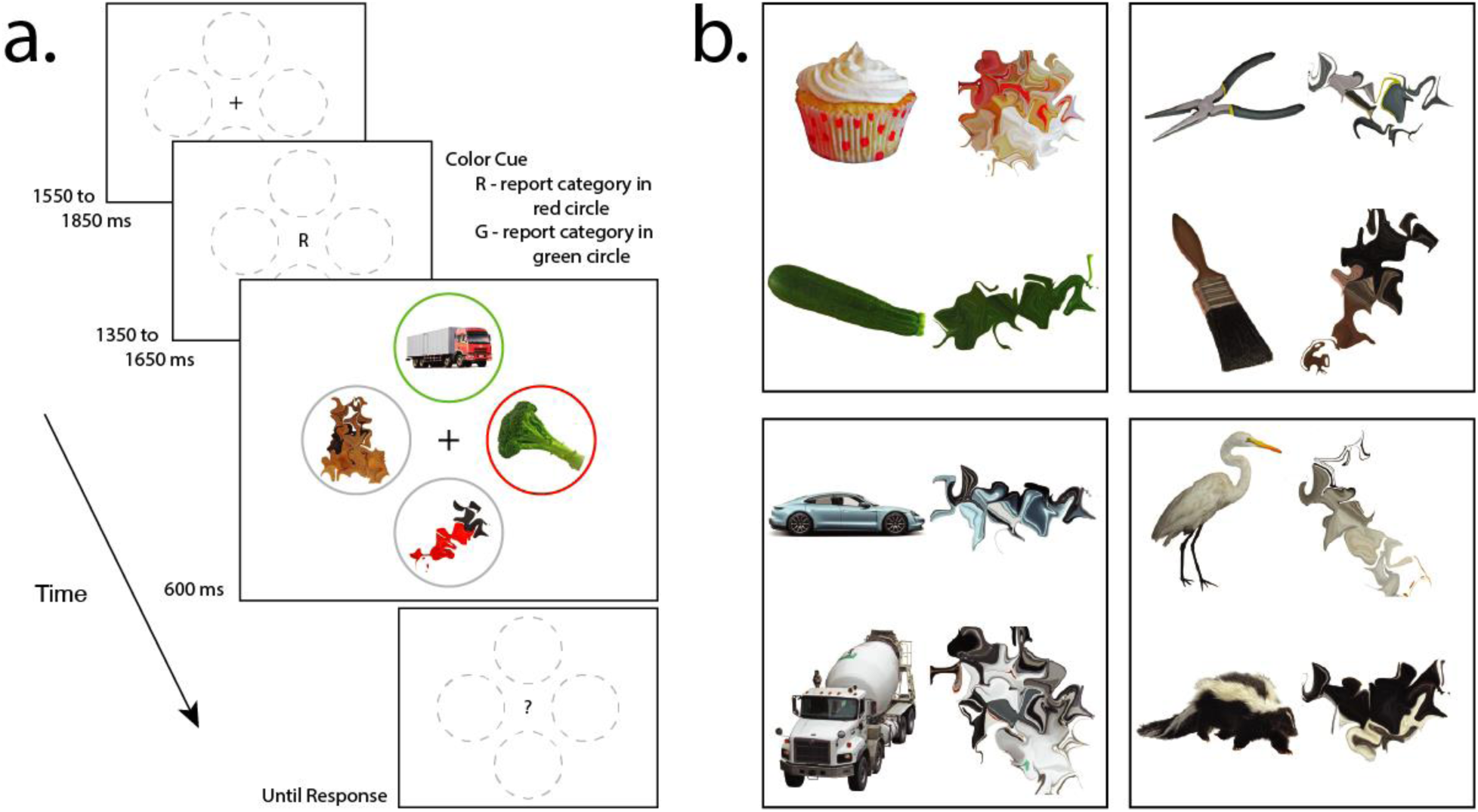
– a.) Trial schematic. Participants reported the category of the object appearing in the cued annulus. b.) Stimuli examples. Each category contained 40 images. Warped version of the image set were employed as unrecognizable non-targets.

Our design provided a second opportunity to explore the functional role of N_T_ and N2pc. We employed naturalistic stimuli in our experiment, presenting 160 examples of visual object categories as targets and recognizable distractors (Figure 1b). Displays always contained one target and one salient distractor and participants were required to report the target category. This provided an opportunity to investigate the role of target identity and distractor identity in the creation of target-elicited and distractor-elicited N_T_. We approached experimentation with the expectation that variance in target identity would create variance in target-elicited N_T_, but were less clear on whether variance in distractor identity would also impact target-elicited N_T_. As the distractor was presented on the vertical in relevant conditions – and thus could not directly create lateralized visual brain activity – covariance between distractor identity and target-elicited N_T_ might imply that the target-elicited N_T_ is sensitive to distractor interference, in line with the idea that the underlying brain activity is involved in resolution of this interference.

We use linear mixed modelling (LMM) in analysis of our results, and this motivates a final, independent goal of the paper: to provide an outline and cookbook for the use of LMM in analysis of multimodal cognitive neuroscience data. Results from psychology and cognitive neuroscience are commonly hierarchical – participants complete trials, and data varies both at the level of participants and at the level of trials. We tend to approach such data by conducting statistical analysis in 2 steps, summarizing results across trials before conducting inferential statistical tests based on the location and distribution of these summarized scores. For example, in repeated-measures analysis of ERP results, we might first calculate per-participant conditional ERPs through mean averaging before conducting a paired t-test to identify a difference between conditional ERPs. Or, in analysis of functional MRI, we might first calculate per-participant beta weights for a voxel before conducting an ANOVA to identify a pattern in the set of beta weights across conditions.

By distinguishing between *fixed effects* and *random effects,* mixed models simplify this analysis. The definition of these terms is surprisingly contentious, but in the current context it is useful to think of fixed effects as those where the sample (nearly) exhausts the underlying population – such as when a stimulus can be a target or distractor in a design and both possibilities are included in the model, or when we sample a continuous variable like RT and have enough data to represent the functional range of the variable reasonably well. Random effects, in contrast, are those where a small part of the population is sampled – as when we employ a few different images as stimuli and these are taken from the population of all images, or when we employ a selection of participants taken from the population of all people (Green & Tukey, 1960).

LMM has a broader scope than 2-step analysis. It provides opportunities to vastly increase statistical power through treatment of trial-wise variance in random effects and is particularly potent in the identification of trial-wise relationships between continuous measures. This type of relationship is of common interest in multimodal cognitive neuroscience, and LMM is therefore a powerful technique to have in one’s analytic toolbox.

## METHOD

### Participants

Forty-nine individuals were recruited from the University of Birmingham community and gave informed consent before completing the experiment. Two were excluded from analysis due to excessive eye movement artefacts in the EEG resulting in rejection of more than 40% of trials, two were excluded as results showed no evidence of the target-elicited N_T_ (ie. the lateral target-elicited signal showed contralateral positive polarity more often than negative polarity; per-participant p < 0.05, binomial test), and a single participant was excluded due to low task accuracy (<70% accuracy). The final sample of 44 individuals had a mean age of 21.9 +/− 2.7 years SD with 23 self-reporting as women, 21 as men, and none as any other gender. Three reported being left-handed. The total duration of the experiment was approximately 2.5 hours and participants were paid £10 per hour. Ethical approval was obtained from the STEM ethics committee at the University of Birmingham and the work adhered to the ethical principles of the Declaration of Helsinki.

### Experimental Task

Trial structure is illustrated in Figure 1a. Participants completed 4 blocks of a cued visual search task and 4 blocks of a repetition detection task in interleaved blocks, with each block composed of 128 trials and the order of tasks counterbalanced across participants. The visual search task took approximately 75% of the experiment duration and only results from this task are treated here.

Each visual search trial began with presentation of a fixation mark on white background (1550 – 1850 ms, randomly selected from uniform distribution) that also contained 4 grey circles located above and below central fixation, and on the left and right (5.7° visual angle). The fixation mark was subsequently replaced by the letter ‘R’ or ‘G’ (1350 – 1650 ms) to indicate the color identifying the target. In the search array, 2 of the grey circles acquired red or green color, and these circles came to contain coherent, recognizable examples of one of 4 visual object categories – tools, vehicles, animals, or foods (Figure 1b). Importantly, in each trial one of either the red or green circles – containing complete objects – appeared on the vertical meridian of the display, with the other presented on the horizontal meridian. Image examples were taken from the BOSS database (Brodeur, Dionne-Dostie, Montreuil, & Lepage, 2010) or from public online image repositories, there were 40 examples in each of the 4 categories, and examples were selected from each category randomly for each trial without replacement across trials until the set was exhausted and the process reset.

Participants had up to 2 s to respond based on the categorical identity of the item in the cued circle, pressing one button on a standard keyboard with their index finger when the target was taken from 2 of the categories and another button with their middle finger when the target was taken from either of the two remaining categories. The mapping of response to category was counterbalanced across participants and the response associated with the target was always opposite to that associated with the distractor object. The two remaining grey circles came to contain unrecognizable morphed versions of the same image set (Figure 1b; Stojanoski & Cusack, 2014; max distortion 160, nsteps 4, iteration 9). The experiment began with verbal instructions and participants then completed training until they achieved ∼80% task accuracy. Stimuli were presented via a 59.5 cm x 33.5 cm LCD monitor at 60 Hz and a distance of 60 cm using PsychToolbox 3 (Brainard, 1997) for Matlab (Mathwords, MA, USA).

### Data acquisition and preprocessing

Electrophysiological data was acquired from sintered Ag/AgCl electrodes at 1 kHz using a Biosemi ActiveTwo amplifier. EEG was collected from 128 electrodes fitted in an elastic cap at equidistant encephalic sites, horizontal electrooculogram (HEOG) was collected from 2 electrodes located 1 cm lateral to the external canthi of the left and right eye, vertical electrooculogram (VEOG) was collected from 2 electrodes immediately above and below the right eye, and unused reference signals were collected from 2 electrodes placed over the left and right mastoid processes. EEG was resampled offline to 200 Hz, digitally filtered with symmetric Hamming windowed finite-impulse response kernels (high-pass at 0.05 Hz, –6 dB at 0.025 Hz; low-pass at 45 Hz, – 6 dB at 50.6 Hz), referenced to the average of encephalic channels, and baselined on the 200 ms interval preceding stimulus onset. Noisy channels were visually identified and interpolated using spherical spline interpolation. Independent component analysis (ICA) of combined EEG and EOG data was used to identify data variance resulting from eye movements. Trials with eye movements in the 500 ms interval following stimulus onset were removed from analysis and the ICA components reflecting eye artefacts were subsequently removed from the data.

### Modelling and statistical inference

Linear mixed modelling relied on fitlme.m and ancillary functions implemented in the statistics toolbox (ver. 12.1) for Matlab R2021a (Mathworks, MA, USA), and on R (R core team, 2012) where detailed. Restricted maximum likelihood was used for variance estimation (unless identified otherwise), Akaike Information Criterion (AIC) was used for model comparison, and ANOVA derivations rely on Satterthwaite approximations of degrees of freedom. Trials resulting in incorrect participant response are discarded from modelling analysis and trials with N_T_ value more than 3 SD from the cross-conditional mean are also excluded from analysis. The data set was ultimately composed of 18640 observations. Preprocessed data and annotated analysis code are available at www.cognitionlab.org.

## RESULTS

### Modelling target– and distractor-elicited Nt

Our experimental design manipulated the laterality of salient, complete objects in the target array such that either the target was presented laterally or the distractor was presented laterally. As stimuli on the vertical meridian do not create a lateralized response, this allowed us to isolate the N_T_ discretely evoked by the target (Figure 2a) or distractor (Woodman & Luck, 1999; Hickey, Di Lollo, & McDonald, 2009). Participants appear to have weakly or occasionally attended to the distractor in our experimental design, resulting in an N_T_ in lateral-distractor trials (Figure 2b). The N_T_ was substantively larger for the target than it was for the recognizable distractor.

**Figure 2.**
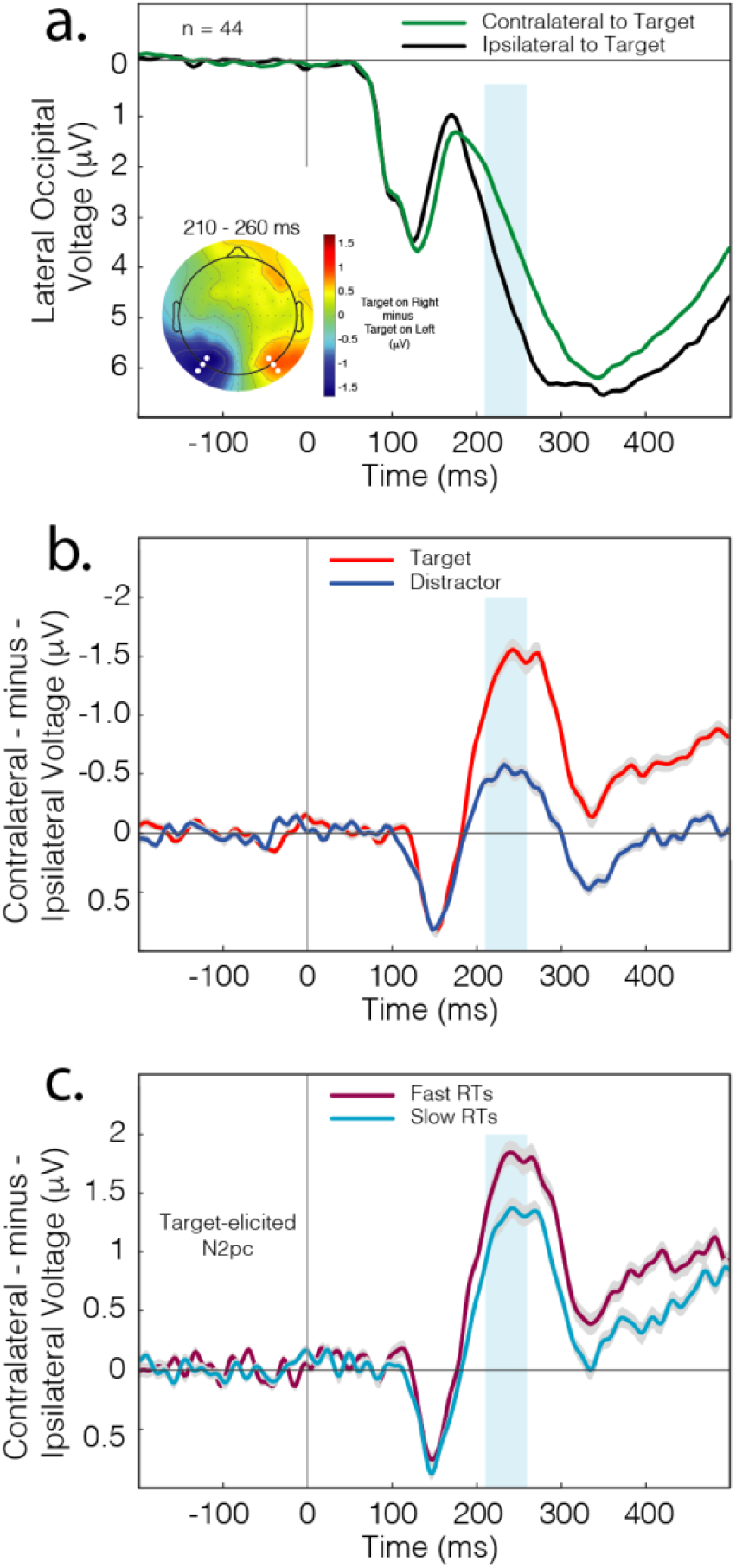
– Experimental results. In panels B and C, error shading reflects bootstrapped SEM. a.) Target-elicited ERP as measured at posterior electrode locations contralateral and ipsilateral to the eliciting stimulus. The topographical map reflects subtraction of right target condition from the left target condition; the N2pc is indicated by blue focus in the left hemisphere and red focus in the right hemisphere. The ERPs in all panels reflect mean signal at the lateral electrodes identified by white marker in the topography. For modelling, the N2pc was measured in the interval identified in blue shading. b.) The target-elicited and distractor-elicited contralateral-minus-ipsilateral difference waves. The N2pc emerges as negative-polarity deviation beginning around 200 ms post-stimulus. c.) The target-elicited N2pc as a function of median split of RT.

Both target-elicited and distractor-elicited N_T_ are preceded by earlier, positive-polarity lateral voltage at about 150 ms that reflects an imbalance in sensory activity between the visual hemispheres. There is a difference in the complexity and spatial frequency of the complete object employed in our stimulus displays, which is presented to one side of the screen, and the unrecognizable degraded object that is presented on the other (Figure 1a). This creates a short-latency lateral effect in the visual ERP, sometimes termed the posterior contralateral positivity (Ppc; Pomerleau et al., 2014) or N1pc (Sänger & Wascher, 2011). This effect is not related to attention and its insensitivity to the behavioural relevance of the evoking stimulus distinguishes it from the subsequent N2pc.

We began analysis by reproducing results from the median split reported by Drisdelle, West, and Jolicoeur (2016). Correct trials were partitioned into ‘Fast RT’ and ‘Slow RT’ subsets for each participant and target-elicited ERPs were calculated for the set of Fast RT subsets and the set of Slow RT subsets separately. As illustrated in Figure 2c, the target-elicited N_T_ preceding fast RTs was larger than the N_T_ preceding slow RTs. As described in the introduction, it is unclear if this difference in N_T_ reflects its impact on information processing and response preparation, rather than shared sensitivity to underlying participant states like arousal or motivation. If it reflects a direct link, the relationship should be discrete to the target-elicited N2pc. If it reflects the influence of participant state, the relationship should emerge for both target-elicited and distractor-elicited N2pc. We used modelling to differentiate between these possibilities. Modelling was based on mean N_T_ amplitude, as measured as mean contralateral-minus-ipsilateral signal over a 50 ms window (210 – 260 ms) centered on the cross-conditional N_T_ peak (235 ms).

Though we were conceptually interested in the idea that N_T_ might predict RT, we modelled N2pc as a function of RT, and this deserves some clarification. RT has a natural lower limit, and there is a higher probability that participants will occasionally make very slow responses rather than very fast ones. As a result, RT has a heavily skewed distribution. While any form of distribution can be modeled with LMMs, a non-normal distribution of the dependent variable often leads to a non-normal distribution of the residuals of the model. This does not invalidate the model as such – it can still be used validly to make predictions – but it introduces challenges and ambiguity in statistical inference. Modelling of RT therefore tends to involve transformation of the dependent variable or the use of generalized linear modelling (Lo & Andrews, 2015).

Both solutions have the downside of making parameter estimates and effect sizes harder to interpret, and of increasing the complexity and computational expense of modelling. In contrast to RT, there is no a priori expectation of non-normal distribution in the Nt, and therefore higher likelihood that module residuals will distribute normally. Our first motivation for modelling N_T_ as a function of RT was therefore to circumvent the need for data transformation or more complex modelling by employing a dependent variable that we expected to distribute normally and a predictor that we expected to express skew. Our second motivation was that modelling of N_T_ allowed us to test hypotheses regarding the sensitivity of N_T_ to variance in target or distractor identity, as outlined in the introduction and described further below.

Our approach to modelling and parameter evaluation involved establishment of a base model with core experimental parameters. We subsequently added random effects that might improve the model, evaluating their efficacy through consideration of AIC. In all cases, models were initiated with a ‘maximal’ parameter structure that included all random effects suggested by the fixed effect structure (Barr, Levy, Scheepers, & Tily, 2013). The random effects were subsequently simplified and the impact of this was assessed using AIC. In Wilkinson notation, the base model was as follows:

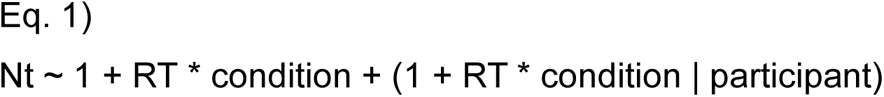

Where N_T_ describes trial N_T_ amplitude, *RT* describes trial reaction time, *condition* describes if the eliciting stimulus was target or distractor, *participant* describes the participant identity, and random effects within levels of *participant* are identified in brackets. Note that for exposition we have made the correlated intercepts explicit in notation for this base model, but do not do so in the remaining equations (except when the intercept is the only factor) as these are absent by default in the notation for most LMM software packages.

We expanded the base model by adding a set of random effects to capture variance created by changes in the target image. This predictor, *target_image,* had levels corresponding to the 160 images that could characterize the target.

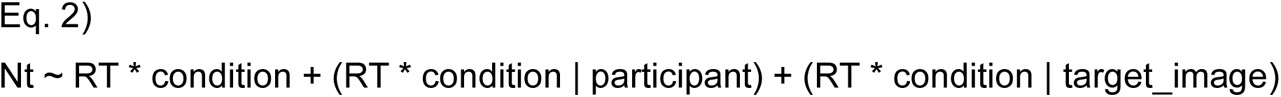

The new random effect was simplified to remove parameters corresponding to the interaction of *RT* and *condition*, then the effect of *RT*, and finally the effect of *condition* to leave only the intercept. Based on AIC, the best of these models included the intercept and the effect of *condition*:

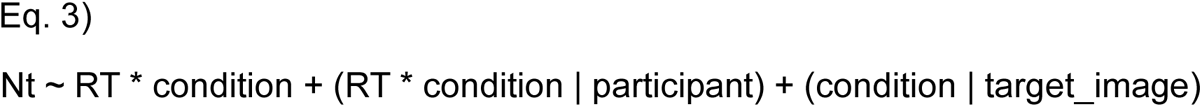

The model in Eq 3 was an improvement over the base model (AIC –104), and we accordingly expanded from this model by adding a second set of random effects to capture variance created by changes in the identity of the distractor image. This predictor, *distractor_image,* had levels corresponding to the 160 images that could characterize the distractor.

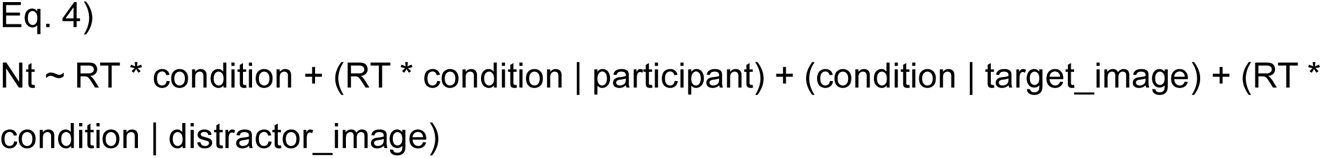

Using the same approach to parameter evaluation, we simplified the structure of the *distractor_image* random effect (AIC –45), and of the *participant* random effect (AIC –20), to identify the final omnibus model.

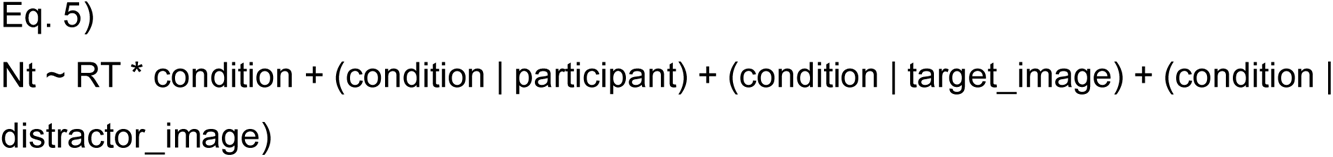

Before drawing inferences from this model we considered if the data met the assumptions of LMM.

### Assumption 1 – Normality of residuals

We visually assessed this in two ways. First, we scatter-plotted the model predictions against model residuals for each of the target and distractor conditions separately and generated empirical probability density functions for these results (Figure 3a). The scatter graph and probability distributions suggest that the N2pc residuals approximate normality with no clear indication of skew or kurtosis. Our second approach was to generate a quartile-quartile (QQ) plot of all N_T_ residuals (collapsed across *condition*; Figure 3b). This identified relatively minor deviation from normality. The LMM is in fact robust to deviation from distributional assumptions (eg. Schielzeth et al., 2020), and we concluded that this deviation was not sufficient to warrant concern.

**Figure 3.**
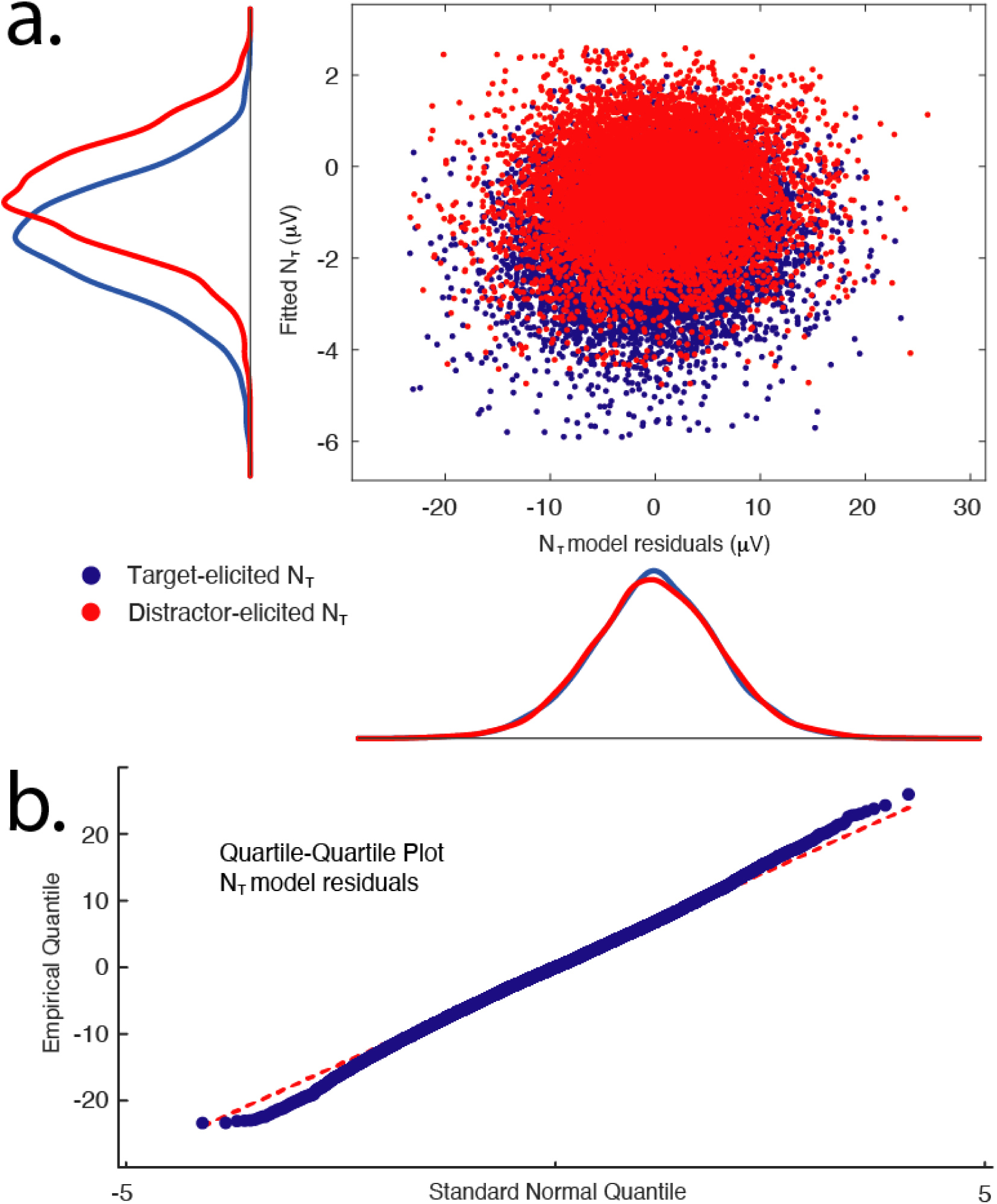
– Evaluation of model assumptions – normality of model residuals. a.) N_T_ model residuals appear normally distributed in both target and distractor conditions. The variance of model residuals does not appear to consistently differ between the target and distractor conditions or as a function of fitted N_T_ amplitude. b.) The QQ plot identifies some minor deviation from normality in the model residuals, but not sufficient to cause concern.

### Assumption 2 – Stability of variance in residuals (homoscedasticity)

LMM assumes that the variance of model residuals is stable across the range of categorical or continuous predictor values. To evaluate this assumption, we scatter-plotted the model residuals for each trial against RT observed in that trial (Figure 4). There is a tear-drop shape to the result that could suggest a reduction in residual variance as RT increases, but careful consideration reveals that this is largely a product of decreasing density of the scatter-plot. That is, the range of residual values at short RT is very much the same as the range of residual values at long RT, but the number of observations decreases. To numerically assess this, we conducted a median split of each participant dataset to isolate the fastest half of trials and the slowest half of trials and calculated the variance of residual values in these data subsets. There was very little difference (28.37 vs 28.10).

**Figure 4.**
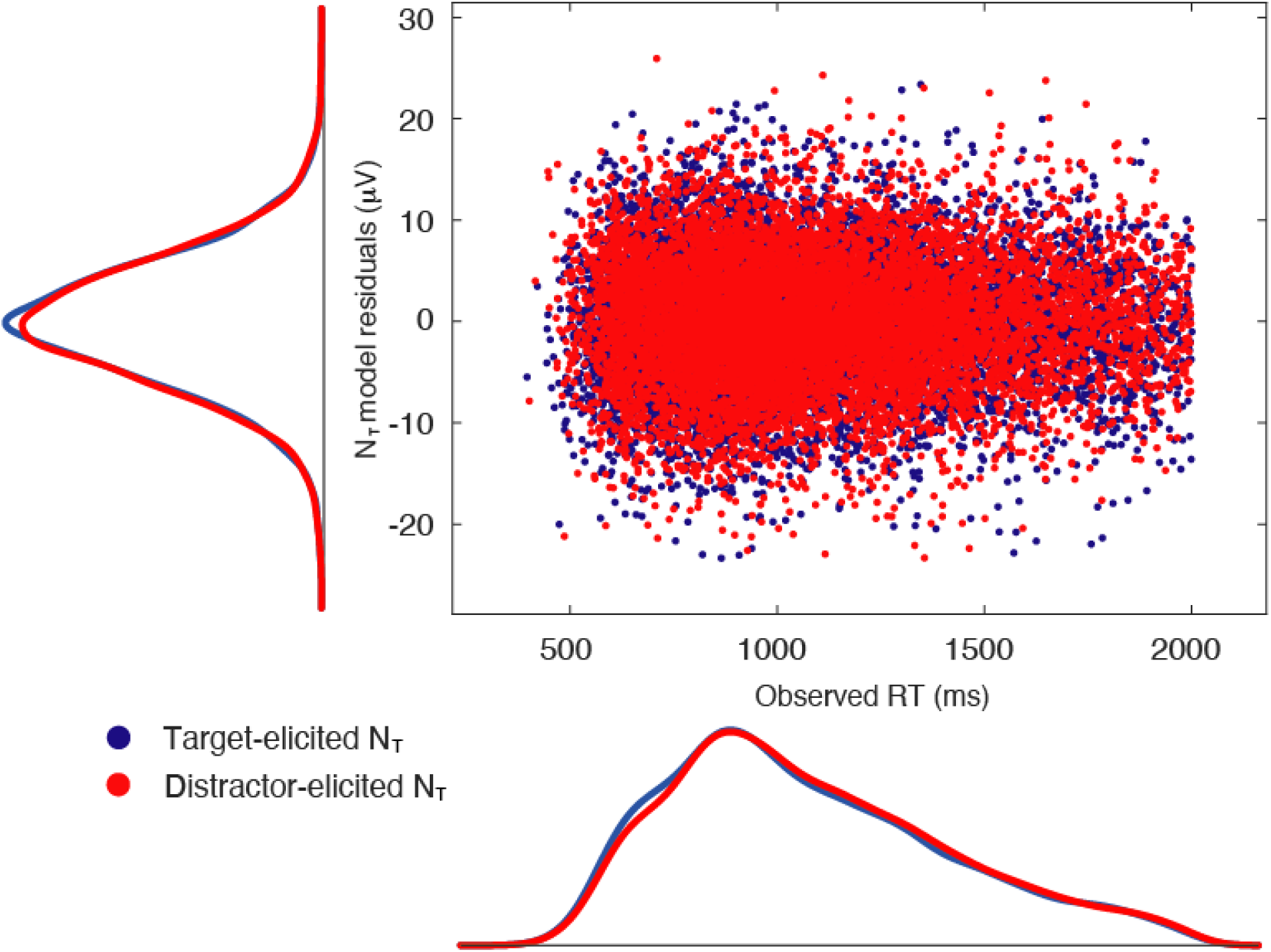
– Evaluation of model assumptions – homoscedasticity. Variance in N_T_ model residuals appears stable across the range of RT predictor values.

### Assumption 3 – Normality of random effects

The normality of random effects is an assumption of LMM but appears to have little influence on fixed effects parameter estimates (Maas & Hox, 2004). However, the impact of deviation from this assumption on the interpretation of the random effect itself has been the subject of less investigation, and, as we draw inferences regarding random effects below, it is reasonable for us to evaluate this assumption. The omnibus model has six random effects: the intercept within levels of *participant,* the intercept and effect of *condition* within levels of *target_image,* and the intercept and effect of *condition* within levels of *distractor_image*. Each of the target and distractor models have 3 random effects: the intercept within levels of *participant,* the intercept within levels of *target_image,* and the intercept within levels of *distractor_image.* Figure 5a illustrates the distribution of random effects for the omnibus model, and Figures 5b and 5c do so for the target and distractor model respectively. As the sample for each of these effects is relatively small – 44 participants, 160 images – the histograms do not describe a precisely normal distribution, but there is no clear evidence of skew or kurtosis and the results appear to approximate the assumption.

**Figure 5.**
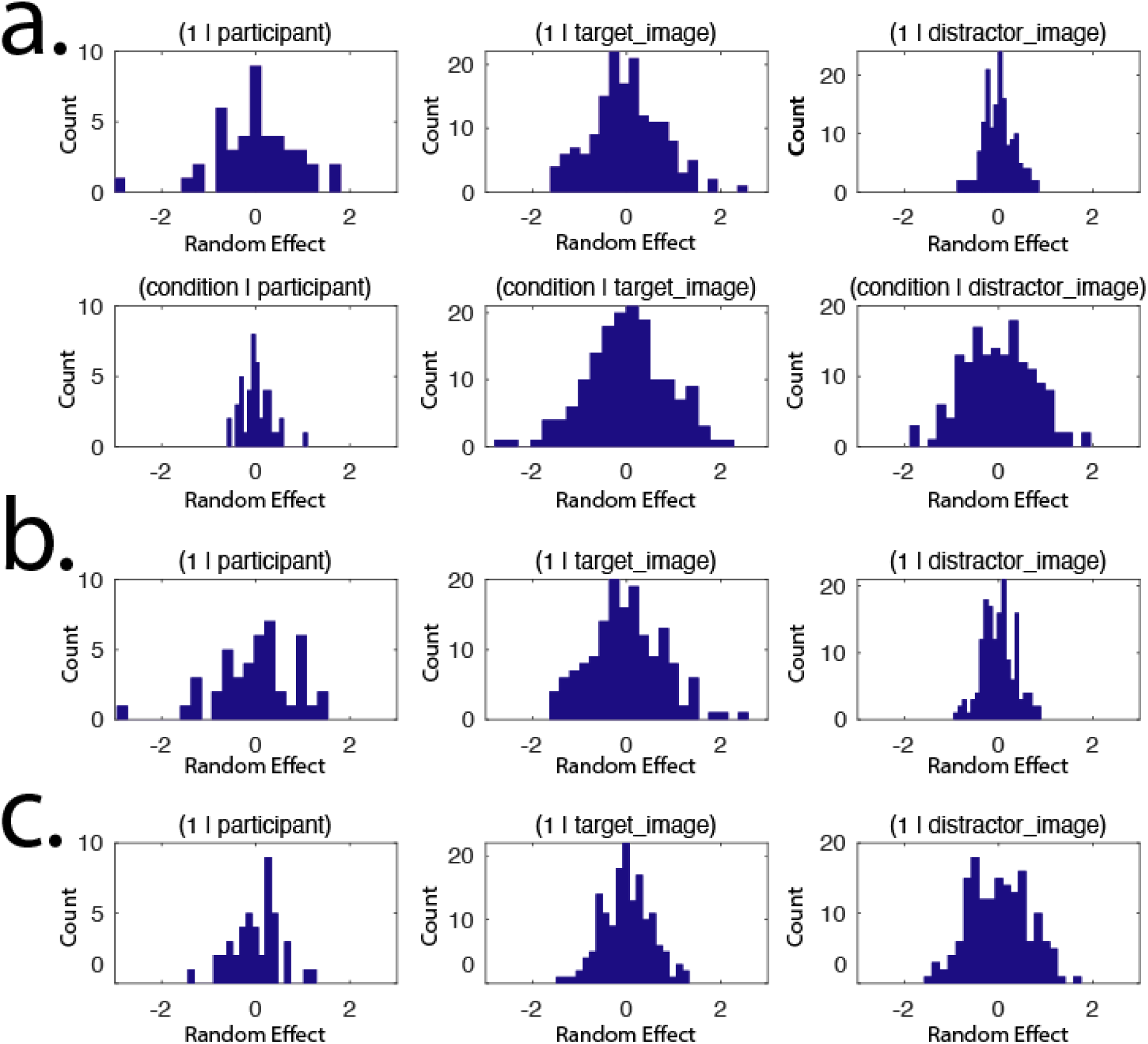
– Evaluation of model assumptions – random effect distributions. a.) Random effects from omnibus model. b.) Random effects from target model. c.) Random effects from distractor model.

### Assumption 4 – Effect linearity

LMM is, of course, unsuited if effects are non-linear. To evaluate this assumption, we scatter-plotted *RT* and N_T_ results separately for each level of *condition* for a subset of participants individually and for all participants at once (Figure 6a). While the raw data is highly variable, there is no immediate evidence of any non-linear pattern in N_T_ as a function of RT. As to be expected, fitted results (Figure 6b) show a clearly linear pattern with reduced variance.

**Figure 6.**
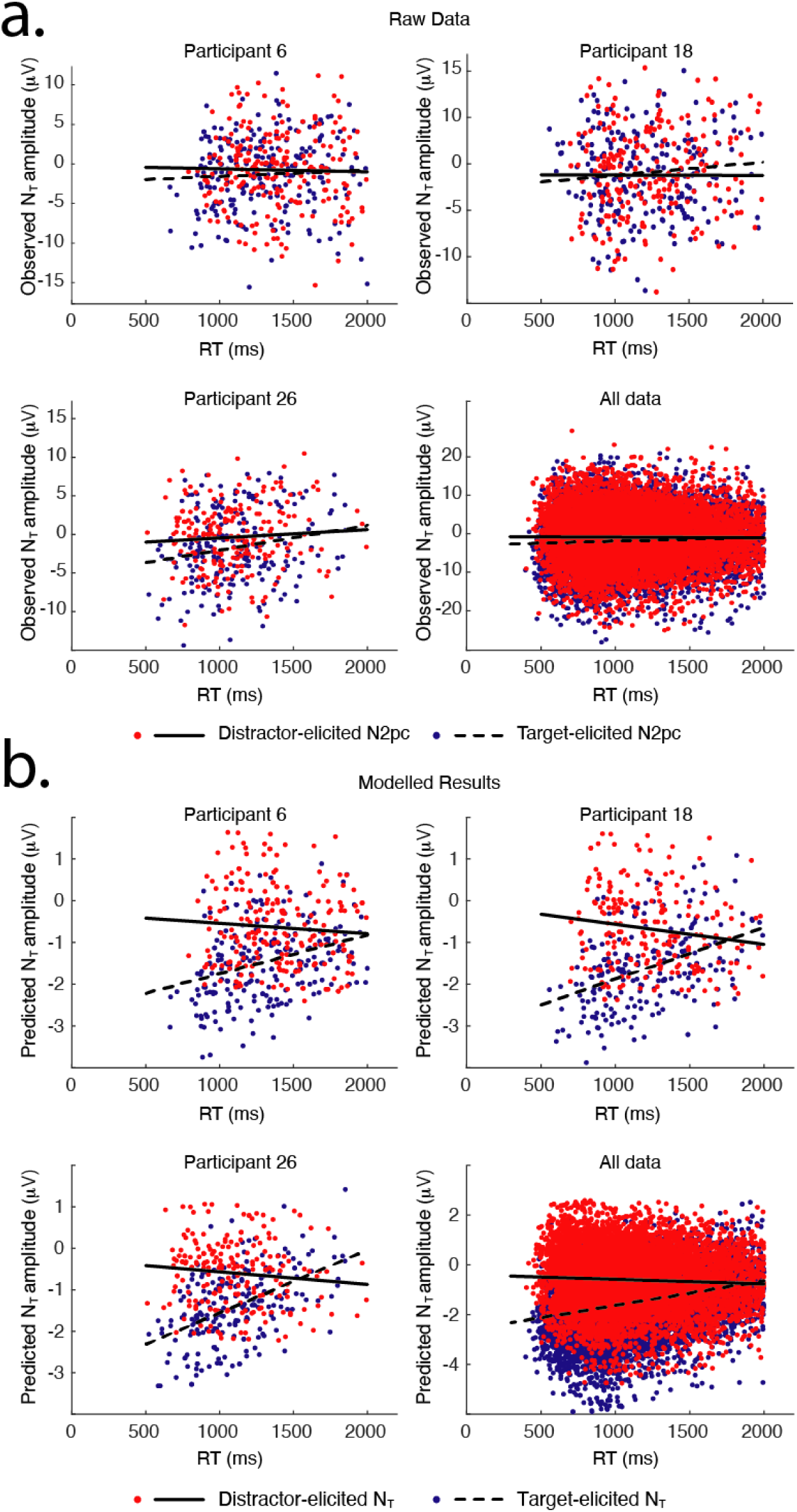
– Evaluation of model assumptions – linearity. a.) Observed N_T_ as a function of observed RT. b.) Modelled N_T_ as a function of observed RT.

### Assumption 5 – Collinearity and multicollinearity

*Collinearity* refers to a problem in linear regression where pairs of predictors are correlated, preventing the model from assigning variance to a unique source. *Multicollinearity* is a slightly broader concept that can emerge in models with 3 or more predictors, where a linear relationship can emerge across the set of predictors (even when pairwise correlations between the predictors do not emerge). Verification of the absence of multicollinearity is therefore more complicated than is the case for collinearity. One approach relies on measurement of *variance inflation factor* (VIF), or on *generalized VIF* (gVIF) when the model contains one or more categorical predictor (Fox & Monette, 1992). Detail on the derivation and meaning of VIF is outside the scope of this paper (Stine, 1995, is a good starting place for further exploration), but the R (R core team, 2021) package *car* (companion to Fox & Weisberg, 2018) includes a function to compute VIF and gVIF. To our knowledge, no equivalent tool is available in the MATLAB environment. There is a substantive caveat to using VIF in this way: in models including interaction terms, VIF and gVIF are inflated and therefore difficult to interpret. VIF values below 3 are generally interpreted as reflecting low multicollinearity; the three terms in our omnibus model, which includes an interaction, hover around this threshold (*condition* gVIF *=* 1.71, *RT* gVIF = 2.79, interaction gVIF = 3.39). We therefore conclude that no further treatment is required.

### Statistical inference

Given that our results appear suitable for LMM analysis, we move on to draw statistical inferences from the models. Our experimental design is such that we have a roughly balanced fixed effect structure – there are similar numbers of trials in each level of *condition* for each participant, and each unique observation of N_T_ is associated with a unique observation of RT. As a result, inferential analysis of fixed effects via ANOVA is accurate and well-suited. This is not strictly necessary – model parameters are inferentially evaluated as part of model building, and there are approaches that allow for direct inference from these values. But the ANOVA is easy, familiar, and comfortable for editors and reviewers (even if its use introduces a second step in analysis, impinging on the grace of single-step mixed modelling).

ANOVA analysis of our omnibus model (Eq 5) with Satterthwaite approximation of degrees of freedom identified a critical interaction of *condition* and *RT* [F(1, 636) = 16.04, p = 6.93×10^−5^]. This also identified an effect of *condition* [F(1,429) = 38.13, p = 1.54×10^−9^] and of *RT* [F(1,3990) = 19.95, p = 8.19×10^−6^]. Note that while the effect of condition is unambiguous, the effect of RT is based on analysis of observations in the reference level of the factor condition, which happened to be when the N_T_ was elicited by the target. This is not a main effect in the sense commonly expected from an ANOVA. The interaction indicates that the effect of RT in the alternative distractor condition differed from the effect of RT in this reference condition. This key result motivates follow up analysis to identify the effect of RT in each of the target and distractor conditions separately.

We followed up to investigate the relationship between N_T_ and RT in target-elicited and distractor-elicited N_T_ conditions separately. Modelling was again completed incrementally, beginning from a base model with addition and simplification of random factors, until the following final model was achieved. Analysis of target-elicited N_T_ identified the following model:

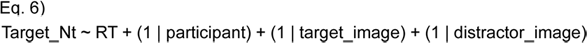

Where *Target_Nt* describes target-elicited N_T_ amplitude.

ANOVA analysis identified a main effect of *RT* on *Target_Nt* [F(1, 5722) = 13.41, p = 2.5 x 10^−4^]. As smaller RT values predicted an a more negative N_T_ amplitude, the parameter estimate for the effect was positive (0.0008 µV increase in negative-polarity N_T_ for every 1 ms increase in the RT). We used the Snijders and Bosker (2011; Correl, Mellinger, & Pedersen, 2021) approach to derive eta squared as a measure of effect size (η^2^ = 0.00091). Note that effect sizes in LMM are substantively smaller than are expected in analysis based on mean averaged data as they reflect the variance explained across all observations in the dataset, rather than the variance explained across a smaller set of averages (which are, by definition, more stable). Rule-of-thumb evaluation of small, medium, and large magnitude effects do not yet exist for LMM, but the reporting of effect size continues to have value for subsequent reproduction and meta-analysis.

Parameter selection for modelling of distractor-elicited N_T_ identified much the same final model:

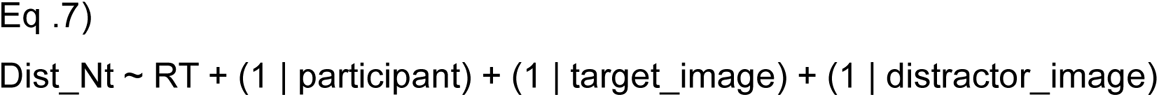

Where *Dist_NT* describes the distractor-elicited N_T_ amplitude.

ANOVA analysis did not identify a reliable effect of *RT* on *Dist_Nt* [F(1, 2499) = 0.04, p = 0.834; η^2^ = –0.05188]. Note that the effect size has negative value. Eta squared can take negative value, and this should be informally interpreted as zero, but provision of the actual variable can forestall bias in reproduction and meta-analysis (Okada, 2016).

### The impact of image variance on target-elicited Nt

Our modelling of target-elicited N_T_ identifies random effects that capture variance as a function of the specific image employed, and inclusion of these effects improved the model. To draw statistical inferences regarding these effects, we used a likelihood ratio test to compare a model containing each random effect to a model identical but for the absence of this effect (Snijders & Bosker, 2012). Importantly, this necessitates a slight change in modelling details: log likelihood comparison of models with a different number of parameters is not valid when variance is estimated using restricted maximum likelihood (REML). For this kind of analysis, maximum likelihood (ML) variance estimation must be employed. We re-calculated the model described in Eq 6 with ML variance estimation and calculated new models missing the random effects of interest.

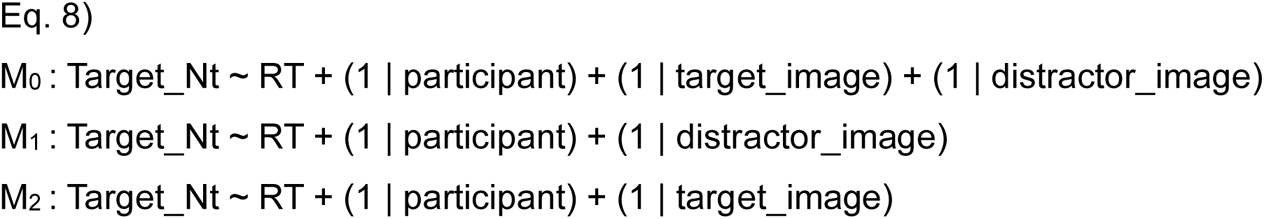

The likelihood ratio statistic distributes as χ² with degrees of freedom corresponding to the difference in the number of parameters between the models. As expected, a contrast of the full model (M_0_) to the model restricted to exclude variance in the target image (M_1_) identified a significant difference (χ²(1) = 74.70, p < 1 x 10^−16^). Variance in the target identity thus played a clear role in determining the amplitude of target-elicited N_T_. More interestingly, a contrast of the full model (M_0_) to the model restricted to exclude variance in the distractor image (M_2_) also identified a significant difference (χ²(1) = 14.15, p = 1.69 x 10^−4^). Variance in the distractor identity thus also played a role in determining the amplitude of the target-elicited N_T_.

Results from analysis of the random effects suggest that the target-elicited N_T_ is sensitive to variance in the distractor image, even though this image is presented on the vertical meridian of the visual field and therefore cannot produce lateralized brain activity itself. We explored this finding in two ways. First, we Pearson-correlated the random intercept for target images with the random intercept with distractor images. This did not identify a relationship (r = –0.071, p = 0.378). The images having greatest impact on the target-elicited N_T_ when serving as targets were therefore not the same images that had the greatest impact on the target-elicited N_T_ when they served as distractor.

We expanded from this by extracting each image from its white background and calculating its mean *luminance*. *Luminance* is an image processing construct that is used in the derivation of a grayscale image from RGB video (as defined by the International Telecommunications Union – Radiocommunications unit, Recommendation BT.601). It is a simple weighting function of RGB values to approximate the perceived luminance of each image pixel, without consideration of physical hardware, viewing circumstance, or psychophysical perceived brightness. In the current context, we calculated the mean *luminance* of each image as an approximation of the perceived brightness of the images for the experimental participants.

We Pearson-correlated the mean *luminance* for each of the 160 images with the random effect for that image, which reflected the modelled impact of the image on the NT. In analysis of the target image, this identified a significant relationship (r = 0.212, p = 0.008). That is, reduced *luminance* of the target image predicted a larger negative-polarity Nt. As the images were presented on a white background, this likely reflects the impact of image contrast and salience: when the image was darker, and therefore of higher contrast with the background, it generated a larger N_T_. However, no such effect emerged in corresponding analysis of the distractor image (r = –0.068, p = 0.402). Increased contrast of the distractor image therefore did not reliably predict a change in the target-elicited N_T_.

## DISCUSSION

Our results demonstrate that target-elicited N_T_ is discretely predicted by RT. This relationship does not emerge for distractor-elicited Nt, suggesting that the link between target-elicited N_T_ and RT is direct and functional, rather than mediated by a third variable such as motivation or arousal. This links N_T_ closely to target processing and is consistent with the notion that mechanisms reflected in N_T_ support the resolution and propagation of target information to downstream cognitive systems, including response preparation and execution. Because of this role in cognition, a larger N_T_ ultimately leads to a quicker response.

We find further that the target-elicited N_T_ is sensitive to variance in the specific images employed in a given trial. That is, we employ 160 images in our design, organized in 4 categories, and the target and recognizable distractor are selected from this set in each trial. Modelling demonstrates that variance in the target image has a reliable impact on the target-elicited N_T_. This is perhaps unsurprising. However, we also find that the target-elicited N_T_ varies as a function of which image is employed as the distractor. As the distractor appears on the vertical meridian of the visual field in relevant conditions, and cannot itself produce lateralized brain activity, this suggests that the distractor identity is in some way interfering with resolution of the target.

One possibility is that this interference is simple distraction. That is, assuming that attention is a fixed resource (eg. Kahneman, 1973), selection of the vertical distractor may reduce participant ability to resolve the target, and therefore reduce the amplitude of target-elicited N_T_. However, the viable alternative is that the relationship between distractor identity and target-elicited N2pc reflects the need for resolution of distractor interference, as suggested by early accounts of N2pc functionality (Luck, et al., 1997). The two accounts make opposite predictions: if it is distraction, then an increase in the ability of the distractor to draw attention – for example, when the distractor is of greater raw visual salience – should presumably result in a decrease of target-elicited N_T_. If it is suppression, then the prediction reverses: as the distractor becomes more interfering, the target-elicited N_T_ – which reflects resolution of this interference – should grow larger.

We attempted to test these predictions in our data by calculating mean *luminance* for each image, as a proxy for object contrast and perceived salience, and using this value to predict variance in target-elicited N_T_ across images. This identified a significant relationship between target-elicited N_T_ and target luminance: higher contrast target images generated a larger amplitude target-elicited N_T_. However, when these same images served as distractor, luminance did not predict target-elicited N_T_ variance. The current results therefore do not discriminate between the distraction and suppression hypotheses above.

There is the opportunity for future work in this context. A more powerful experimental design might parametrically manipulate distractor salience – perhaps in more than one way – such that the effect on target-elicited N_T_ could be sampled across a large range of distractor salience. One challenge to this venture would be to operationally define distractor salience. Here, we have adopted a very simple definition: raw *luminance* contrast with the background. A more advanced approach might use statistical models of salience (eg. Itti & Koch, 2001), or characterize the image salience experimentally in an orthogonal task. There is, moreover, the possibility that the relationship between distractor identity and target-elicited N_T_ that is identified in the current data is not a simple product of physical salience but reflects interference at the level of shared visual features, or shared semantics. These are more challenging hypotheses to test, as they would require modelling of the interaction of target and distractor identity, and this would require vastly more experimental data.

The current results thus demonstrate that the target-elicited N_T_, and therefore N2pc, is closely involved in target processing (Eimer, 1996), and plays a critical role in the sheltering and propagation of target information to downstream cognition. However, our results leave open the possibility that this mechanism of target sheltering acts in whole or part through the resolution of distractor interference, in line with early interpretation of the component (eg. Luck et al., 1997).

A second goal for the current paper was to demonstrate the efficacy of LMM in analysis of multimodal cognitive neuroscience data. We used LMM to show a trial-wise relationship between an ERP component and manual RT and show how the concurrent modelling of random effects can vastly increase statistical power to detect such a relationship. We moreover show that a random effect can be interesting and interpretable in of itself, and that random effect structure can be examined in order to provide insight.

The general analytic principle presented here abstracts to other multimodal data. In other work, we have used LMM to identify trial-wise relationships between phase-locked ERP components and measures of induced oscillatory EEG activity (van Zoest, Huber Huber, Weaver, & Hickey, 2021), between the N2pc and machine-learning classification of EEG data (Acunzo, Grignolio, & Hickey, under review), and to model N2pc amplitude as a function of decision-making variables like information entropy, expected value, and regret (Kobayashi, Barbaro, Gottlieb, & Hickey, under review). LMM is well suited and statistically optimal in any situation where trial-wise variance between measures is of interest.

## Acknowledgments

Our thanks to Holly Ahmed for support and discussion. All authors were supported by the European Research Council (ERC) under the European Union Horizon 2020 Research and Innovation Program (Grant Agreement 804360 to CH).

## Competing interest statement

The authors have no competing interests to declare. Declaration regarding generative AI: No generative AI was used in the preparation of this manuscript.

## Notes

### Competing Interest Statement

The authors have declared no competing interest.

